# Resolving phylogenetic and biochemical barriers to functional expression of heterologous iron-sulphur cluster enzymes

**DOI:** 10.1101/2021.02.02.429153

**Authors:** Helena Shomar, Pierre Simon Garcia, Elena Fernández-Fueyo, Francesca D’Angelo, Martin Pelosse, Rita Rebelo Manuel, Ferhat Büke, Siyi Liu, Niels van den Broek, Nicolas Duraffourg, Carol de Ram, Martin Pabst, Simonetta Gribaldo, Beatrice Py, Sandrine Ollagnier de Choudens, Gregory Bokinsky, Frédéric Barras

## Abstract

Many of the most promising applications of synthetic biology, including engineering of microbes for renewable chemical production, relies upon the ability of genetically-tractable hosts to express heterologous enzymes from foreign species. While countless methods for facilitating heterologous enzyme expression have been developed, comparable tools for facilitating heterologous enzyme activity are generally lacking. Such tools are needed to fully exploit the biosynthetic potential of the natural world. Here, using the model bacterium *Escherichia coli*, we investigate why iron-sulphur (Fe-S) enzymes are often inactive when heterologously expressed. By applying a simple growth complementation assay with collections of Fe-S enzyme orthologs from a wide range of prokaryotic diversity, we uncover a striking correlation between phylogenetic distance and probability of functional expression. Moreover, co-expression of a heterologous Fe-S biogenesis pathway increases the phylogenetic range of orthologs that can be functionally expressed. On the other hand, we find that heterologous Fe-S enzymes that require specific electron carrier proteins within their natural host are rarely functionally expressed unless their specific reducing partners are identified and co-expressed. We demonstrate *in vitro* that such selectivity in part derives from a need for low-potential electron donors. Our results clarify how phylogenetic distance and electron transfer biochemistry each separately impact functional heterologous expression and provide insight into how these barriers can be overcome for successful microbial engineering involving Fe-S enzymes.

## Introduction

Microbes acquire new phenotypes *via* horizontal gene transfer, a process that reshapes microbial evolution and ecology. Gene transfer is also an everyday laboratory technique that enables expression of heterologous proteins and engineered biosynthetic pathways. In natural systems, obstacles against foreign gene expression, which include restriction-modification systems and differences in codon usage, can impede horizontal gene transfer from influencing host phenotype. In the lab, such genetic obstacles are routinely avoided to ensure heterologous protein expression, *e.g.* by using strains that lack genome defence systems or by optimizing codon usage. However, heterologous proteins must not only be expressed but also must retain their activity to affect the host phenotype or to function as part of an engineered pathway. Unlike obstacles to heterologous protein expression, the obstacles that prevent heterologous protein activity are rarely studied^1^. One enzyme class that is particularly prone to inactivity in heterologous hosts is iron-sulphur cluster (Fe-S) enzymes^2–5^. Fe-S enzymes use bound iron and sulphur metalloclusters to catalyse a variety of essential reactions, including many that are relevant to biotechnology such as nitrogen reduction, dehydration, sulphur insertion, and methyl transfer^6^. Identifying obstacles to functional heterologous expression of Fe-S proteins is therefore an important challenge.

Fe-S enzymes are maturated by dedicated Fe-S biogenesis pathways and proteins **(Figure 1A)**. If a heterologous Fe-S enzyme cannot obtain Fe-S clusters from its host, enzyme activity is impaired. How Fe-S enzymes obtain clusters from Fe-S biogenesis proteins continues to be intensively investigated in both prokaryotic and eukaryotic models^7,8^. Two Fe-S biogenesis pathways, ISC and SUF, catalyse assembly and delivery of Fe-S clusters to many apo-target proteins in prokaryotes (*e.g.* more than 150 in *Escherichia coli*). Common to both ISC and SUF is the use of cysteine desulphurase enzymes (IscS or SufSE) to obtain sulphur from L-cysteine and scaffold proteins to assemble Fe-S clusters (IscU or SufBCD). Fe-S cluster carrier proteins of the A-type (IscA, SufA, ErpA) Nfu-type (NfuA) or ApbC-type (Mrp) subsequently deliver clusters to apo-targets^9,10^. Moreover, some Fe-S require additional factors to sustain catalytic turnover, particularly electron carrier proteins^11^. This requirement may present an additional barrier to heterologous expression.

**Figure 1.**
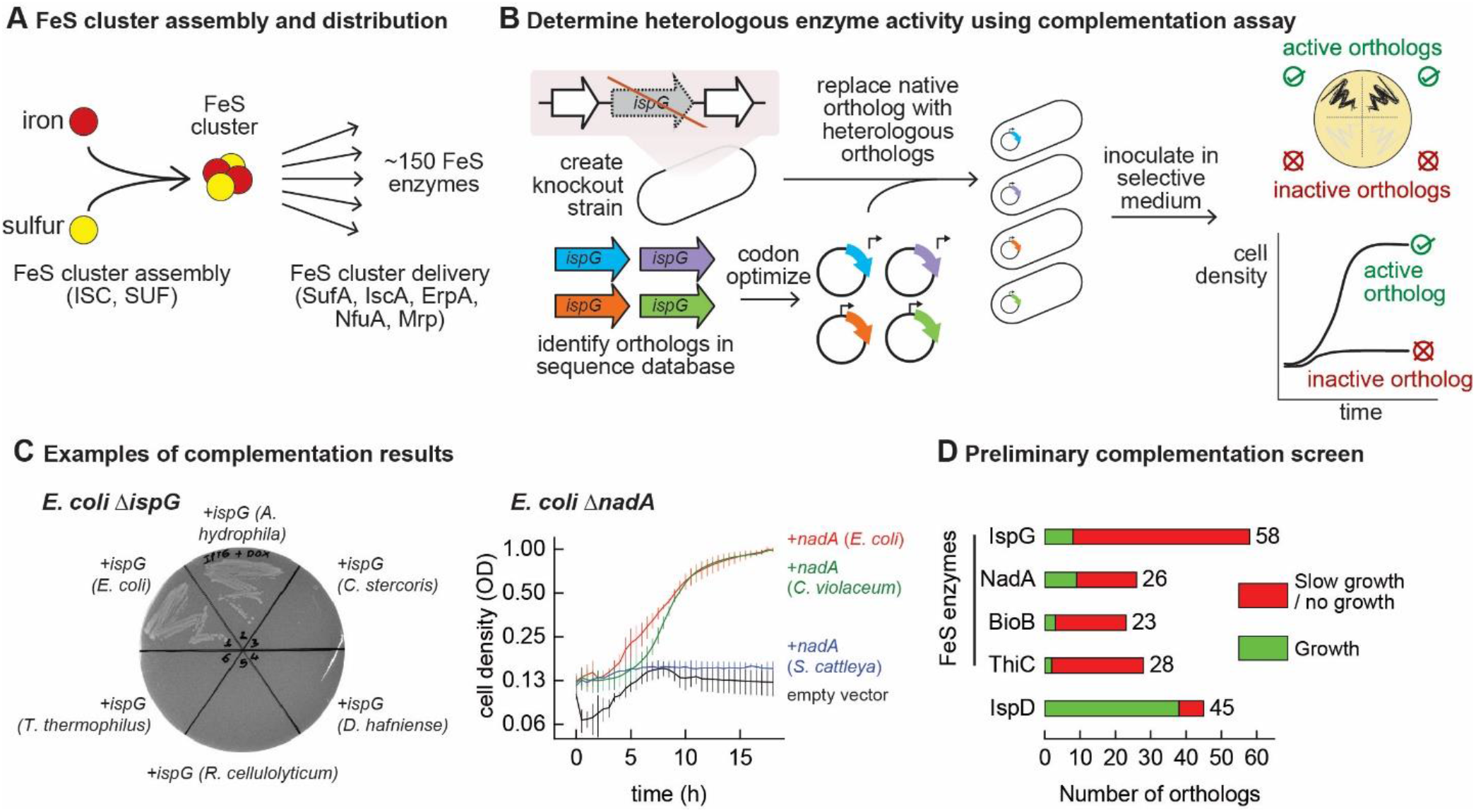
Mapping functional expression of heterologous Fe-S enzymes using complementation experiments. **A.** In prokaryotes, two pathways that assemble and deliver Fe-S clusters have been identified (ISC, SUF), as have several Fe-S delivery proteins (SufA, IscA, ErpA, NfuA in *E. coli*). **B.** The complementation assays used to determine activity of heterologous orthologs expressed by *E. coli.* Orthologs of conditionally-essential enzymes are identified in sequence databases, codon-optimized, and cloned into a low-copy plasmid. Expression is controlled by an inducible promoter (P_Tet_). The plasmids are then transformed into *E. coli* strains lacking the corresponding *E. coli* ortholog. Each strain is inoculated in selective media and expression of the heterologous ortholog is induced. Growth in selective media indicates that the heterologous ortholog retains sufficient activity to support growth of *E. coli.* **C.** Exemplary results of complementation assay obtained using either solid or liquid selective medium. Error bars represent standard error of duplicate cultures inoculated from separate colonies. **D.** Results of preliminary complementation tests comparing activity of Fe-S enzymes with a non-Fe-S enzyme. Orthologs tested for the preliminary screen are listed in Table S1.

The full biotechnological potential of Fe-S enzymes cannot be harnessed until we understand why many of them lose activity when heterologously expressed. A lack of systematic data prevents any meaningful prediction on whether a heterologous Fe-S enzyme will remain active within a host. Defining the limits of horizontal transfer of Fe-S enzymes between species is therefore needed to identify the specific factors that impede functional heterologous expression. Here, we combine bioinformatics with an *in vivo* platform to map the transferability of several biotechnologically-interesting Fe-S enzymes between prokaryotes. We also characterize at a molecular level the requirements for sustaining the heterologous activity of an Fe-S enzyme from the radical *S-*adenosylmethionine (rSAM) methyltransferase family.

## Results

### Heterologous Fe-S enzymes are less likely to be functionally expressed than non-Fe-S enzymes

We performed a preliminary screen to evaluate the ability of Fe-S enzymes to retain activity when expressed in a heterologous host. We constructed four strains of *E. coli* MG1655 that each lack a conditionally-essential Fe-S enzyme: NadA (quinolinate synthase), IspG (4-hydroxy-3-methylbut-2-enyl-diphosphate synthase), BioB (biotin synthase), and ThiC (4-amino-5-hydroxymethyl-2-methylpyrimidine phosphate synthase). These proteins were chosen because they are involved in metabolic pathways for valuable compounds (*e.g.* vitamins, biofuels, and fragrances). Orthologs of these enzymes were identified in genomes belonging to a diverse range of bacterial phyla (*e.g. Actinobacteria, Firmicutes, Proteobacteria, Cyanobacteria*) for a preliminary survey **(Table S1)**. Each orthologous gene was codon-optimized, cloned into a low-copy expression vector, and transformed into the corresponding *E. coli* knockout strain. The activities of the heterologous enzymes were tested using a complementation assay: growth in selective media indicates that the heterologous enzyme is functional when expressed in *E. coli* **(Figure 1B, 1C)**. The majority of Fe-S enzyme orthologs (115 out of 135) failed to complement growth **(Figure 1D, Table S1).** For comparison, we tested orthologs of a non-Fe-S enzyme (IspD, 2-C-methyl-D-erythritol 4-phosphate cytidylyltransferase) for functional expression in *E. coli*. In contrast with the Fe-S enzymes tested, the majority of IspD orthologs (38 out of 45) complemented growth of *E. coli* Δ*ispD,* including many orthologs from species whose IspG orthologs failed to complement growth of *E. coli* Δ*ispG* **(Table S1)**. Of note, average sequence identity percentage of the heterologous IspG and IspD orthologs with the *E. coli* orthologs were 45% and 36%, respectively. Altogether, these observations suggest that Fe-S enzymes are far more likely to lose activity within a heterologous host as compared to enzymes whose activities do not require Fe-S cluster or other post-translational modifications.

### Correlating compatibility of heterologous enzymes with phylogenetic distance from host ortholog

We next sought to explore in detail the factors that determine whether a Fe-S enzyme retains its activity within a heterologous host. We hypothesized that heterologous Fe-S enzymes whose sequences are less similar to the host *E. coli* ortholog might be less likely to complement growth. To test this hypothesis, we focused on orthologs of NadA and IspG. We first built a protein database that is representative of known prokaryotic diversity, comprising 248 prokaryotic (Archaea and Bacteria) proteomes corresponding to 48 phyla, 96 classes, and 218 orders **(Table S2)**. This database contains 65 proteomes that were used for preliminary screen. We identified 192 and 196 orthologs of NadA and IspG, respectively **(Table S3)**, which were used to infer a Maximum Likelihood phylogeny **(Figure 2A, D)**. Next, we used a clustering approach based on phylogenetic distances to select representative sequences to be experimentally tested, which yielded 47 sequences for NadA and IspG each **(Tables S3, S4)**. The distribution of patristic distances of the selected orthologs from the *E. coli* NadA and IspG sequences closely matched with the patristic distance distribution obtained from all enzyme homologs, indicating that the selected orthologs are representative of the entire database of known orthologs **(Figure 2B, E)**. Furthermore, a a two-sided Wilcoxon test between full and sampled distances from *E. coli* rejected the hypothesis of significant differences between the selected set of orthologs (47 both) and the set of all known orthologs (191 for NadA and 195 for IspG, *p*-value = 0.4714 for NadA and *p*-value = 0.4739 for IspG). Altogether, these results indicate that the set of orthologs tested experimentally is a representative sample of the phylogenetic diversity of NadA and IspG sequences across prokaryotes.

**Figure 2.**
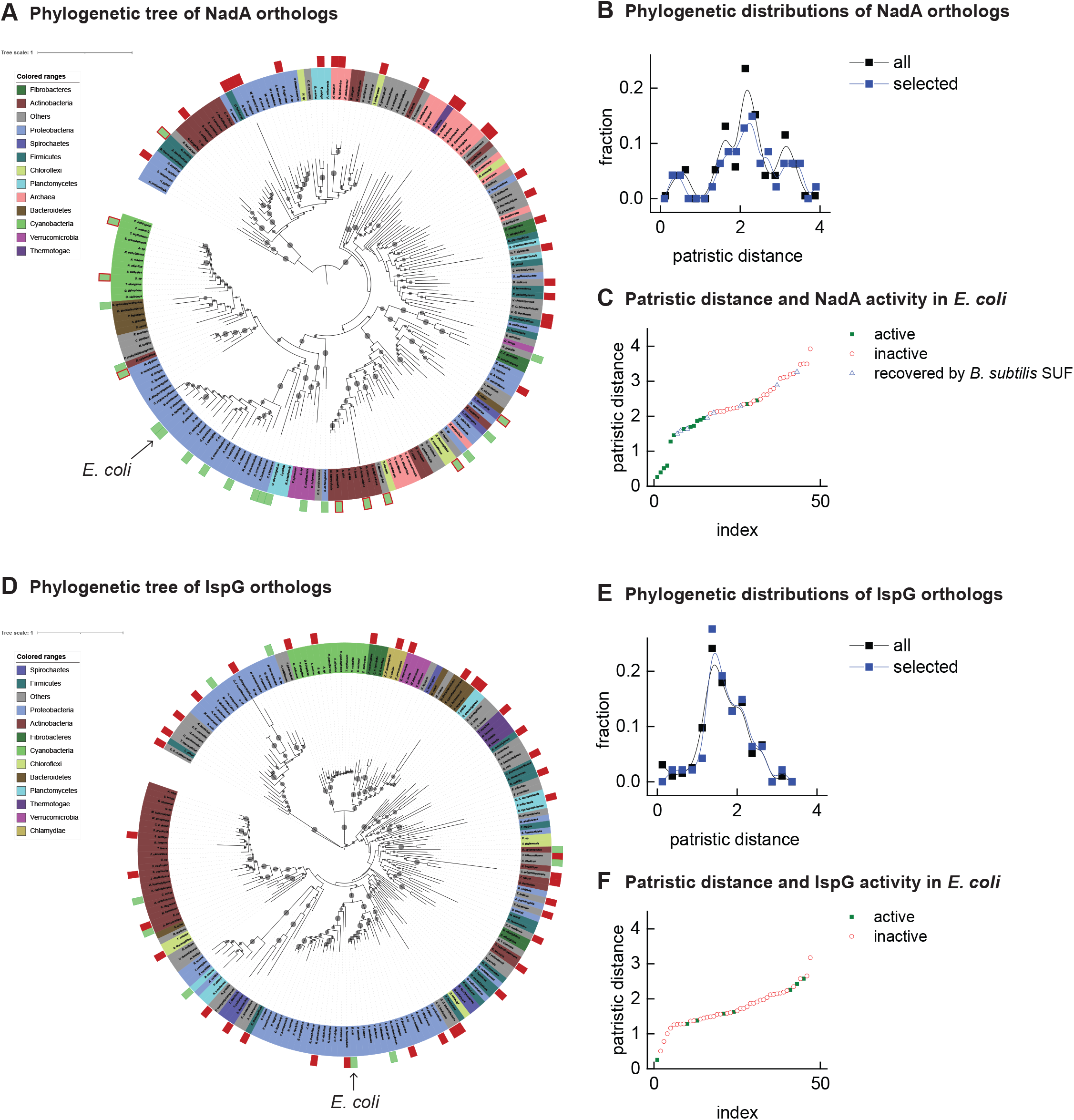
Phylogenetic maps contrasting the compatibility of Fe-S enzymes NadA **(A-C)** and IspG **(D-F)** across a selection of orthologs representative of known prokaryotic sequence diversity. **A, D:** Phylogeny of NadA orthologs **(A)** and IspG orthologs **(D)**. In each diagram, the position of *E. coli* ortholog is highlighted by an arrow. Orthologs selected for our complementation assay are indicated by color rectangles (green: successful complementation; green surrounded by red: complementation by co-expression of *B. subtilis* SUF; red: no complementation) **B, E:** Distributions of patristic distances from the corresponding *E. coli* ortholog of all sequences (black curves) and the sequences of orthologs selected for testing (blue curves)**. C, F:** Complementation and reactivation results correlated with patristic distance from the *E. coli* ortholog.

Sequences from our 47 phylogenetically-representative sets of IspG and NadA were each then codon-optimized for expression in *E. coli* and tested using our complementation assay. Among the NadA orthologs representative of phylogenetic diversity, 14 out of 47 orthologs complemented growth of *E. coli* Δ*nadA,* indicating functional expression **(Table 1)**. Consistent with our hypothesis, NadA orthologs that were functionally expressed in *E. coli* exhibited a low patristic distance from the *E. coli* NadA ortholog **(Figure 2C)**. The high correlation factor (*η*^2^) between heterologous expression and patristic distance (*η*^2^ = 0.596) suggests that phylogenetic proximity is a useful predictor of whether a NadA ortholog can be functionally expressed in *E. coli*.

**Table 1.** Results of NadA complementation of *E. coli* Δ*nadA*. Minus (−): negative complementation; plus (+): positive complementation; N.D.: not determined.

| Phylum | Species/Strain | Accession number | Complementation |  |  |
| --- | --- | --- | --- | --- | --- |
|  |  |  | NadA | NadA + *B. subtilis* SUF | NadA + *E. coli* SUF |
| Actinobacteria | *Mycolicibacterium smegmatis* MC2 155 | ABK69796.1 | - | + | + |
|  | *Rubrobacter xylanophilus* DSM 9941 | ABG03084.1 | + | N.D. | N.D. |
|  | *Collinsella stercoris* DSM 13279 | EEA90923.1 | - | - | - |
|  | *Acidimicrobium ferrooxidans* DSM 10331 | ACU53899.1 | - | + | - |
|  | *Streptomyces cattleya* NRRL 8057 = DSM 46488 | AEW93658.1 | - | - | - |
| Bacteroidetes | *Salinivirga cyanobacteriivorans* | ALO15705.1 | + | N.D. | N.D. |
| Candidatus Fermentibacteria | *Candidatus Fermentibacter daniensis* | KZD19396.1 | - | - | - |
| Candidatus Melainabacteria | *Candidatus Gastranaerophilales bacterium HUM_7* | DAA95005.1 | - | - | - |
| Chloroflexi | *Sphaerobacter thermophilus* DSM 20745 | ACZ38300.1 | - | + | - |
|  | *Thermogemmatispora tikiterensis* | RAQ94418.1 | - | - | - |
| Chrysiogenetes | *Desulfurispirillum indicum* S5 | ADU65000.1 | - | - | - |
| Cyanobacteria | *Synechocystis* sp. PCC 6803 | BAA18685.1 | - | + | - |
|  | *Crocospaera watsonii* WH 8501 | WP_007304059.1 | - | + | - |
| Deinococcus-Thermus | *Thermus thermophilus* HB27 | AAS80968.1 | - | + | - |
| Dictyoglomi | *Dictyoglomus thermophilum* H-6-12 | ACI18692.1 | - | - | - |
| Euryarchaeota | *Methanopyrus kandleri* AV19 | AAM01290.1 | - | - | - |
|  | *Haloferax volcanii* DS2 | ADE02976.1 | - | - | - |
|  | *Haloterrigena turkmenica* DSM 5511 | ADB59428.1 | - | - | - |
|  | *Candidatus Methanomethylophilus alvus* Mx1201 | AGI85733.1 | - | - | - |
| Fibrobacteres | *Chitinivibrio alkaliphilus* ACh1 | ERP31080.1 | - | - | - |
| Firmicutes | *Bacillus subtilis* subsp. *subtilis* str. 168 | CAB14745.1 | - | + | + |
|  | *Desulfitobacterium hafniense* DCB-2 | ACL18980.1 | - | - | - |
|  | *Ruminiclostridium cellulolyticum* H10 | ACL77768.1 | - | - | - |
|  | *Helioibacterium modesticaldum* Ice1 | ABZ83886.1 | - | - | - |
| Nitrospinae | *Nitrospina gracilis* | CCQ89850.1 | + | N.D. | N.D. |
| Planctomycetes | *Gemmata obscuriglobus* | WP_010044270.1 | + | N.D. | N.D. |
|  | *Pirellula staleyi* DSM 6068 | ADB17048.1 | - | - | - |
| Proteobacteria | *Aeromonas hydrophila* | ABK37682.1 | + | N.D. | N.D. |
|  | *Desulfovibrio vulgaris* str. *Hildenborough* | AAS96285.1 | - | - | - |
|  | *Allochromatium vinosum* | ADC62035.1 | + | N.D. | N.D. |
|  | *Arcobacter butzleri* | WP_020848486.1 | - | - | - |
|  | *Novosphingobium stygium* | SCY25206.1 | - | + | + |
|  | *Herminiimonas arsenicoxydans* | CAL61268.1 | + | N.D. | N.D. |
|  | *Chromobacterium violaceum* ATCC 12472 | AAQ61340.1 | + | N.D. | N.D. |
|  | *Bdellovibrio bacteriovorus* HD100 | CAE81099.1 | + | N.D. | N.D. |
|  | *Methylobacillus flagellatus* KT | ABE50646.1 | - | - | - |
|  | *Anaeromyxobacter dehalogenans* 2CP-C | ABC83113.1 | + | N.D. | N.D. |
|  | *Candidatus Pelagibacter ubique* HTCC1002 | EAS84697.1 | - | - | - |
|  | *Syntrophobacter fumaroxidans* MPOB | ABK17287.1 | - | - | - |
|  | *Cellvibrio japonicus* Ueda107 | ACE83018.1 | + | N.D. | N.D. |
|  | *Escherichia coli* str. K-12 substr. MG1655 | AAC73837.1 | + | N.D. | N.D. |
|  | *Desulfarculus baarsii* DSM 2075 | ADK83407.1 | - | - | - |
|  | *Salinisphaera* sp. LB1 | AWN14904.1 | + | N.D. | N.D. |
| *Spirochaetes* | *Leptospira interrogans* serovar Lai str. 56601 | AAN47817.2 | + | N.D. | N.D. |
|  | *Spirochaeta thermophila* DSM 6578 | AEJ62405.1 | - | + | - |
| *Synergistetes* | *Thermanaerovibrio acidaminovorans* DSM 6589 | ACZ19971.1 | - | - | - |
| *Thermotogae* | *Thermotoga maritima* MSB8 | AAD36711.1 | - | - | - |
| *Verrucomicrobia* | *Coralimargarita akajimensis* DSM 45221 | ADE55989.1 | + | N.D. | N.D. |

Among the 47 IspG orthologs selected from a representative set, only 8 complemented growth of *E. coli* Δ*ispG* **(Table 2)**. The sequences of functionally-expressed IspG orthologs did not group together in the IspG phylogeny but were instead patchily dispersed across the entire phylogenetic distribution of sequences **(Figure 2F)**. Accordingly, no correlation between functional expression and patristic distance from *E. coli* was observed (*η*^2^ = 0.003). Altogether, these results indicate that the ability of heterologous IspG orthologs to maintain activity in *E. coli* does not correlate with phylogenetic proximity, in contrast with NadA orthologs.

**Table 2.** Results of the IspG complementation of *E. coli* Δ*ispG*. Minus (-): negative complementation; plus (+): positive complementation; N.D.: not determined.

| Phylum | Species/Strain | Accession number | Complementation |  |  |  |  |
| --- | --- | --- | --- | --- | --- | --- | --- |
|  |  |  | IspG | IspG + *Synechocystis* PetF | IspG + *B. subtilis* YkuP/YkuN | IspG + *S. cattleya* Fdx | IspG + *R. sphaeroides* Fdx |
| *Acidobacteria* | *Candidatus Koribacter versatilis* Ellin345 | ABF40425.1 | - | N.D. | N.D. | N.D. | N.D. |
| *Actinobacteria* | *Thermoleophilum album* | SEH14556.1 | - | N.D. | N.D. | N.D. | N.D. |
|  | *Streptomyces coelicolor* A3 | CAB91132.1 | - | - | - | - | N.D. |
|  | *Rubrobacter xylanophilus* DSM 9941 | ABG04368.1 | + | N.D. | N.D. | N.D. | N.D. |
|  | *Cutibacterium acnes* KPA171202 | AAT83253.1 | + | N.D. | N.D. | N.D. | N.D. |
|  | *Collinsella stercoris* DSM 13279 | EEA90248.1 | - | - | - | - | N.D. |
|  | *Acidimicrobium ferrooxidans* DSM 10331 | ACU53587.1 | + | N.D. | N.D. | N.D. | N.D. |
|  | *Streptomyces cattleya* NRRL 8057 = DSM 46488 | AEW97312.1 | - | - | - | + | N.D. |
|  | *Egibacter* sp. | PWFH01000026.1_54 | - | N.D. | N.D. | N.D. | N.D. |
| *Aquificae* | *Aquifex aeolicus* VF5 | AAC07467.1 | + | N.D. | N.D. | N.D. | N.D. |
|  | *Thermovibrio ammonificans* HB-1 | ADU96577.1 | - | N.D. | N.D. | N.D. | N.D. |
| *Bacteroidetes* | *Pedobacter heparinus* DSM 2366 | ACU05649.1 | - | N.D. | N.D. | N.D. | N.D. |
|  | *Salinivirga cyanobacteriivorans* | ALO15331.1 | - | N.D. | N.D. | N.D. | N.D. |
| *Candidatus Fermentibacteria* | *Candidatus Fermentibacter daniensis* | KZD16972.1 | - | N.D. | N.D. | N.D. | N.D. |
| *Candidatus Melainabacteria* | *Candidatus Caenarcanum bioreactoricola* | MOYB01000001.1_3 | - | N.D. | N.D. | N.D. | N.D. |
|  | *Candidatus Gastranaerophilales bacterium HUM_7* | DAA93923.1 | - | N.D. | N.D. | N.D. | N.D. |
| *Candidatus Sumerlaeota* | *Candidatus Sumerlaea chitinovorans* | AXA34785.1 | - | N.D. | N.D. | N.D. | N.D. |
| *Chlamydiae* | *Chlamydia caviae* GPIC | AAP05169.1 | - | - | - | - | N.D. |
| *Chloroflexi* | *Sphaerobacter thermophilus* DSM 20745 | ACZ37901.1 | - | N.D. | N.D. | N.D. | N.D. |
| *Cyanobacteria* | *Gloeobacter violaceus* PCC 7421 | BAC91563.1 | - | - | N.D. | N.D. | N.D. |
|  | *Synechocystis* sp. PCC 6803 | BAA17717.1 | - | + | - | - | N.D. |
| *Deinococcus-Thermus* | *Thermus thermophilus* HB27 | AAS82019.1 | - | - | - | - | N.D. |
| *Fibrobacteres* | *Fibrobacter succinogenes* subsp. *succinogenes* S85 | ACX75977.1 | - | N.D. | N.D. | N.D. | N.D. |
| *Firmicutes* | *Bacillus subtilis* subsp. *subtilis* str. 168 | CAB14437.1 | - | + | + | + | N.D. |
|  | *Desulfotobacterium hafniense* DCB-2 | ACL21716.1 | - | - | - | - | N.D. |
|  | *Symbiobacterium thermophilum* IAM 14863 | BAD40486.1 | - | N.D. | - | N.D. | N.D. |
|  | *Ruminiclostridium cellulosyticum* H10 | ACL74833.1 | - | - | - | - | N.D. |
| *Lentisphaerae* | *Victivallales bacterium* CCUG 44730 | AVM43904.1 | - | N.D. | N.D. | N.D. | N.D. |
| *Nitrospinae* | *Nitrospina gracilis* | CCQ91393.1 | - | - | N.D. | N.D. | N.D. |
| *Planctomycetes* | *Blastopirellula marina* DSM 3645 | EAQ77076.1 | +/- | + | + | - | N.D. |
|  | *Isosphaera pallida* ATCC 43644 | ADV63232.1 | - | - | - | - | N.D. |
|  | *Phycisphaera mikurensis* NBRC 102666 | BAM02738.1 | - | N.D. | N.D. | N.D. | N.D. |
| Proteobacteria | *Aeromonas hydrophila* | ABK39186.1 | + | N.D. | N.D. | N.D. | N.D. |
|  | *Allochromatium vinosum* | ADC62931.1 | - | N.D. | N.D. | N.D. | N.D. |
|  | *Azoarcus sp. BH72* | CAL93544.1 | + | N.D. | N.D. | N.D. | N.D. |
|  | *Helicobacter pylori J99* | WP_000892444.1 | - | - | - | N.D. | N.D. |
|  | *Methylococcus capsulatus str. Bath* | AAU91397.1 | + | N.D. | N.D. | N.D. | N.D. |
|  | *Rhodobacter sphaeroides 2.4.1* | ABA79143.1 | - | - | - | - | + |
|  | *Mariprofundus ferrooxydans* | EAU54713.1 | - | N.D. | N.D. | N.D. | N.D. |
|  | *Cellvibrio japonicus Ueda107* | ACE82628.1 | - | - | - | - | N.D. |
|  | *Escherichia coli str. K-12 substr. MG1655* | AAC75568.1 | + | N.D. | N.D. | N.D. | N.D. |
|  | *Anaplasma phagocytophilum str. CRT38* | EOA62006.1 | - | N.D. | N.D. | N.D. | - |
| *Spirochaetes* | *Spirochaeta thermophila DSM 6578* | WP_014624852.1 | - | N.D. | N.D. | - | N.D. |
| *Synergistetes* | *Thermanaerovibrio acidaminovorans DSM 6589* | ACZ19163.1 | - | N.D. | N.D. | N.D. | N.D. |
| *Tenericutes* | *Spiroplasma citri* | APE74616.1 | - | N.D. | N.D. | N.D. | N.D. |
| *Thermodesulfobacteria* | *Thermodesulfatator indicus DSM 15286* | AEH44631.1 | - | N.D. | N.D. | N.D. | N.D. |
| *Thermotogae* | *Thermotoga maritima MSB8* | AAD35972.1 | - | - | - | - | N.D. |
| *Verrucomicrobia* | *Coralimargarita akajimensis DSM 45221* | ADE56011.1 | - | - | - | - | N.D. |

We sought to confirm that the inactive NadA and IspG orthologs used for the representative distribution were expressed in our knockout strains. Because even some of the active orthologs could not be detected by SDS-PAGE **(Table S5)**, we used a mass spectroscopy-based shotgun proteomics approach to determine whether 7 inactive NadA orthologs and the 16 inactive IspG orthologs could be detected when expressed in permissive media. We detected all 7 of the inactive NadA orthologs and 10 of the 16 inactive IspG orthologs **(Table S5)**. Given that *E. coli* IspG is largely insoluble when overexpressed^12^, and that insoluble proteins are often difficult to detect using mass spectroscopy, we suspect that insolubility may contribute to inactivation of heterologous IspG orthologs and potentially other Fe-S enzymes. Nevertheless, it is very unlikely that a correlation between phylogenetic proximity and heterologous activity is obscured by poor expression of IspG orthologs.

### Activity of NadA orthologs depends upon Fe-S biogenesis pathway components

We next tested approaches for restoring function of the inactive NadA orthologs. Our finding that the activity of heterologous NadA orthologs correlates with phylogenetic proximity to *E. coli* NadA suggests that co-expression of a heterologous Fe-S pathway might re-activate the heterologous NadA orthologs. We therefore cloned a heterologous Fe-S pathway (the *Bacillus subtilis* SUF operon) into a separate vector (pBbA5a), which was introduced into *E. coli* Δ*nadA* carrying *B. subtilis* NadA **(Table S6)**. Satisfyingly, co-expression of the *B. subtilis* SUF system together with *B. subtilis* NadA fully recovered growth of *E. coli* Δ*nadA* **(Figure S1, Table 1)**. Expression of *B. subtilis* SUF also activated 8 out of 33 additional NadA orthologs **(Table 1)**. By comparison, overexpression of the *E. coli suf* operon activated only 3 of the 8 NadA orthologs activated by *B. subtilis* SUF (*B. subtilis, Mycolicibacterium smegmatis,* and *Novosphingobium stygium*) **(Table 1)**. Thus, use of a heterologous Fe-S pathway may often broaden host compatibility with certain heterologous Fe-S enzymes.

### Compatibility of IspG orthologs is limited by need for taxa-specific electron transfer proteins

While co-expression of either *B. subtilis* or *E. coli* SUF pathways did not recover any inactive IspG ortholog tested, several heterologous IspG orthologs that did not complement *E. coli* Δ*ispG* were surprisingly able to obtain Fe-S clusters *in vitro* when incubated with the *E. coli* Fe-S transfer protein ErpA **(Figure S2)**. These observations suggest that the inactivity of IspG orthologs may be unrelated to Fe-S transfer. The Fe-S cluster of *E. coli* IspG requires electron inputs from a electron carrier protein, flavodoxin 1 (FldA), to sustain catalysis^11^. Although many enzymes that require electron carrier proteins are able to obtain electrons from a variety of carrier proteins^13,14^, the activity of *E. coli* IspG cannot be supported by *E. coli* flavodoxin 2 (FldB)^15^. Therefore we suspected that the inactivity of heterologous IspG orthologs reflects a requirement for a specific electron carrier protein from the original hosts. Specific electron carriers that support activity of *Synechocystis* and *B. subtilis* IspG orthologs have been previously identified. For example, the IspG ortholog from the cyanobacterium *Synechocystis* is known to be reduced by the ferredoxin PetF^16^. We therefore cloned PetF into a pBbA5a vector and co-transformed into *E. coli* Δ*ispG* together with a vector bearing *Synechocystis* IspG. Co-expression of PetF with *Synechocystis* IspG indeed complemented growth of *E. coli* Δ*ispG* **(Table 2)**. Similarly, co-expression of *B. subtilis* flavodoxins observed to activate *B. subtilis* IspG, YkuP and YkuN^17^, allowed complementation of *E. coli* Δ*ispG.*

The specific electron carriers needed to activate the remaining IspG orthologs tested are not known. Furthermore, while prokaryotic genomes usually encode multiple electron carrier proteins, no reliable bioinformatics method for identifying the carrier proteins able to support IspG activity currently exists. Therefore, we first simultaneously co-expressed multiple electron carrier proteins taken from 27 additional microorganisms together with their cognate IspG orthologs. Of the 27 inactive IspG orthologs tested, 6 were re-activated by cognate electron carriers identified within their original host **(Table S7)**. We individually tested the ability of electron carrier proteins from *Rhodobacter sphaeroides, Synechocystis,* and *Streptomyces cattleya* to support the activity of their corresponding IspG ortholog. Strikingly, only 1 of the 5 electron carriers tested from *R. sphaeroides* could activate *R. sphaeroides* IspG. IspG orthologs from *Synechocystis* and *S. cattleya* IspG were also selective for specific carriers taken from their cognate hosts **(Table S6)**.

We next explored the cross-species compatibility of IspG orthologs with heterologous electron transfer proteins obtained from different species **(Table 2 and S7)**. *Synechocystis* PetF activated nearly every *Cyanobacteria* IspG ortholog tested but only a small number of IspG orthologs outside the *Cyanobacteria* phylum. The individual ferredoxins from *S. cattleya* and *R. sphaeroides* further activated additional IspG orthologs from the *Actinobacteria* and *Proteobacteria*, respectively, but very few IspG orthologs outside their respective taxa. Altogether, 19 out of 53 IspG orthologs tested were re-activated by co-expression of either cognate or heterologous electron transfer proteins. Collectively, these results indicate that the requirement for specific electron transfer proteins can prevent functional expression of heterologous IspG.

### A specific double-cluster ferredoxin enhances the activity of the Fe-S-dependent methyltransferase TsrM

The molecular basis underlying the selectivity of Fe-S enzymes for specific electron transfer proteins is poorly understood. We therefore explored the electron transfer protein selectivity of a biotechnologically-relevant Fe-S enzyme^18^. rSAM methyltransferase enzymes rely upon Fe-S clusters to catalyse biosynthesis of valuable natural products, including antibiotics^19,20^. The cobalamin-dependent rSAM methyltransferase TsrM, an enzyme from the thiostrepton synthesis pathway of *Streptomyces laurentii*, uses its cobalamin cofactor as an intermediate carrier to transfer a methyl group to C2 of tryptophan^21^. Although its catalytic cycle does not require an electron donor for each turnover event, TsrM requires one electron to reduce its cobalamin cofactor to cob(I)alamin, which reacts with SAM to form MeCbl **(Figure 3A)**. TsrM can also be inactivated by adventitious oxidation of the bound cobalamin factor from cob(I)alamin to cob(II)alamin. Recovering the catalytically-active cob(I)alamin form requires reduction by an electron transfer protein^22^.

**Figure 3.**
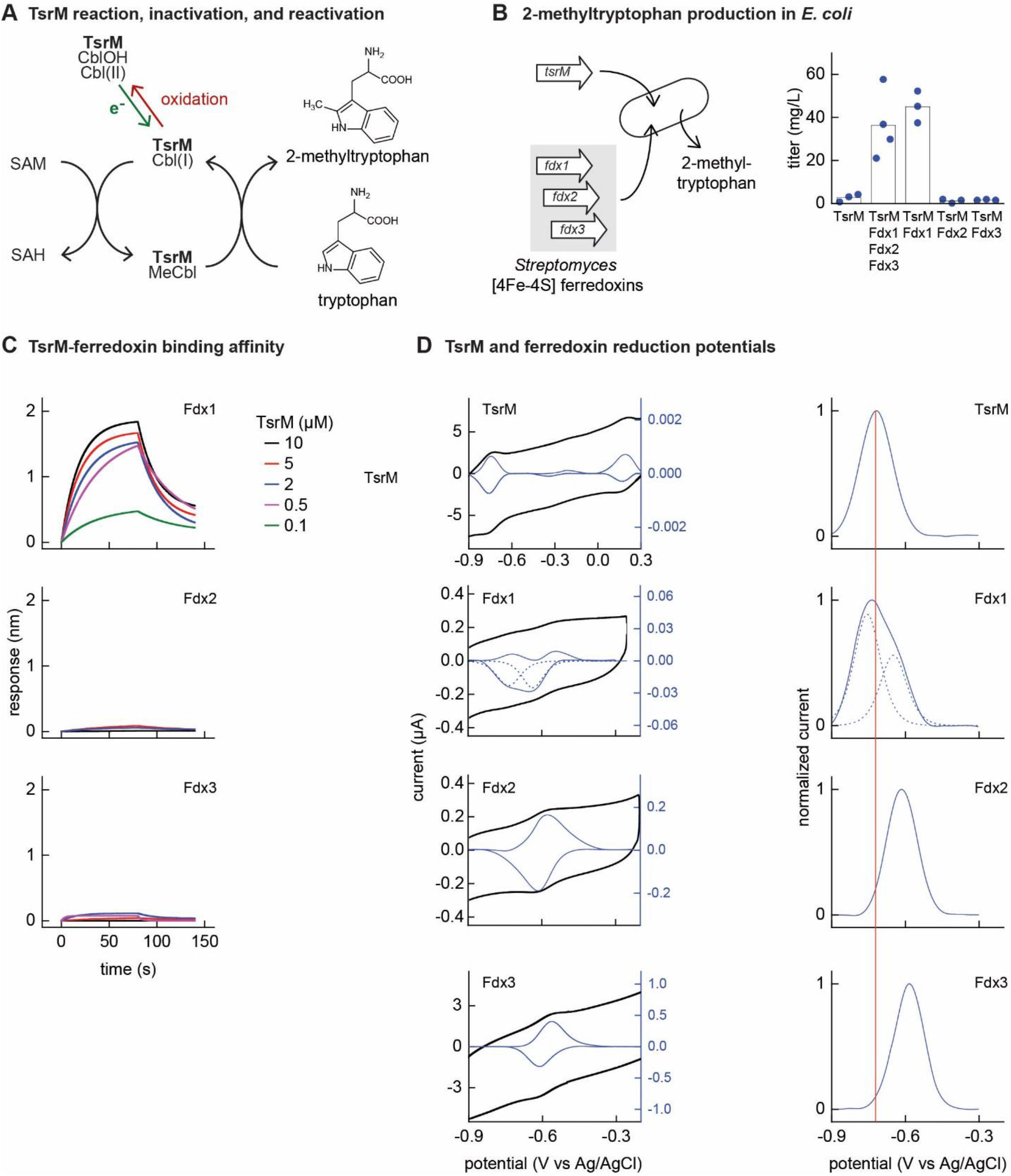
Identifying the barrier to optimal activity of an Fe-S-dependent rSAM enzyme. **A.** The enzyme TsrM synthesizes 2-methyltryptophan as part of the synthesis pathway of the antibiotic thiostrepton. A one-electron reduction (in green) converts the bound cobalamin to cob(I)alamin, which is converted to MeCbl via reaction with SAM. TsrM can be inactivated during catalysis by adventitious oxidation of the bound cob(I)alamin to cob(II)alamin (in red). Recovering the catalytically-active cob(I)alamin form requires a one-electron reduction (green). **B.** Titers of 2-methyltryptophan in a TsrM-expressing *E. coli* strain increase substantially upon co-expression of a specific *S. cattleya* ferredoxin (Fdx1) whereas co-expression of alternative *S. cattleya* [4Fe-4S] ferredoxins, Fdx2 and Fdx3, do not increase titers. **C, D.** Identifying the molecular basis enabling Fdx1 to recover TsrM activity. **C.** Bio-layer interferometry (BLI) of TsrM with Fdx1, Fdx2, and Fdx3 indicates that TsrM interacts measurably with Fdx1 alone. **D.** Cyclic voltammetry of TsrM and the three ferredoxins indicates that the reducing potentials of one of the [Fe-S] clusters of Fdx1 is sufficiently low-potential to reduce TsrM metal cofactors (cob(II)alamin to cob(I)alamin and [4Fe-4S]^+2^ to [4Fe-4S]^+1^) and thus restore activity after adventitious oxidation.

*E. coli* NCM3722 co-expressing *S. laurentii* TsrM together with the *E. coli* cobalamin import pathway (*btu* operon expressed from a plasmid^23^) produced 3 ±2 mg/L 2-methyltryptophan **(Figure 3B)**. We sought to determine whether *in vivo* TsrM activity could be improved using *Streptomyces* [4Fe-4S] ferredoxins, as [4Fe-4S] ferredoxins exhibit low reducing potentials (−700 – −300 mV)^24^ that are expected to match the low reduction potential of cob(II)alamin/cob(I)alamin (< −500 mV vs. NHE)^25,26^. We used three [4Fe-4S] ferredoxins (here designated Fdx1, Fdx2, and Fdx3), two of which (Fdx1 and Fdx3) were able to activate *S. cattleya* IspG **(Table S6)**. Simultaneous co-expression of all three [4Fe-4S] ferredoxins together with TsrM increased 2-methyltryptophan titers to 36 ±16 mg/L. By co-expressing each ferredoxin individually with TsrM, we determined that Fdx1 alone improves activity of TsrM (45 ±7 mg/L 2-methyltryptophan), while expression of Fdx2 and Fdx3 do not increase 2-methyltryptophan titers **(Figure 3B)**.

We further investigated the molecular basis of ferredoxin selectivity by TsrM using *in vitro* experiments with purified proteins. Consistent with previous observations, purified TsrM contains 0.9 cobalamin and 3.5 Fe per monomer **(Figure S3)**. The iron content of the ferredoxins after anaerobic purification or reconstitution with iron and sulphur salts (7.4, 2.9, and 2.8 Fe atoms *per* monomer for Fdx1, Fdx2, and Fdx3 respectively) is consistent with their corresponding annotations as double-cluster and single-cluster ferredoxins. We first used bio-layer interferometry (BLI) to determine whether TsrM associates preferentially with Fdx1. While we were able to detect association between Fdx1 and TsrM (dissociation constant *K*_D_ = 40 μM), we found that Fdx2 and Fdx3 did not measurably associate with TsrM **(Figure 3C)**. We also compared the redox potentials of each ferredoxin against TsrM using voltammetry. The reduction potentials determined for the [4Fe-4S] cluster of TsrM matches with cob(II)alamin/cob(I)alamin (−730 mV vs Ag/AgCl), in agreement with previous reports^15^. The voltammogram of Fdx1 indicated two redox transitions associated with each [4Fe-4S] cluster (−750 mV and −650 mV mV vs Ag/AgCl), the lower of which supports reduction of TsrM metallic cofactors [[4Fe-4S]]^2+^ to [[4Fe-4S]]^1+^ cluster) and (cob(II)alamin to cob(I)alamin). The redox potentials of 4Fe-4S Fdx2 and [4Fe-4S] Fdx3 are each too high for favourable reduction of TsrM metallic cofactors (−620 mV and −585 mV respectively vs Ag/AgCl) **(Figure 3D)**. Therefore, both binding specificity and reduction potential determine productive interactions between electron transport proteins and TsrM, which thus constrains its functional heterologous expression. We note that the TsrM-Fdx1 dissociation constant is consistent with a transient interaction characteristic of an electron transfer process.

## Discussion

Our study has used Fe-S enzymes to map the interspecies versatility of Fe-S biogenesis pathways. In doing so, we have distinguished specific barriers that limit functional transfer of heterologous Fe-S enzymes. Most strikingly, we found that Fe-S pathways are highly compatible with certain heterologous Fe-S enzymes, except at large phylogenetic distances from the host. This is demonstrated by the activity of nearly one third of the heterologous NadA orthologs tested and is also supported by our observation that many IspG orthologs and the rSAM enzyme TsrM require only a specific electron transfer protein to achieve activity in *E. coli*. The versatility of Fe-S pathways is also illustrated by our finding that many of the NadA orthologs taken from organisms that lack an ISC pathway are activated even by the endogenous *E. coli* ISC expressed in our conditions. This versatility, which is likely a consequence of the need for Fe-S pathways to provide Fe-S clusters to many diverse Fe-S enzymes present in the cell, also appears to facilitate functional gene transfer across moderate phylogenetic distances.

While approximately one-third of the NadA orthologs complemented *E. coli* Δ*nadA* and were thus compatible with *E. coli* ISC, several NadA orthologs exhibited activity only when co-expressed with either *E. coli* or *B. subtilis* SUF pathways. Interestingly, several of the NadA orthologs were selectively activated by *B. subtilis* SUF. This may be due to structural differences between components of the *E. coli* and *B. subtilis* SUF systems that effectively restrict Fe-S cluster transfer to orthologs that are closely-related or otherwise similar^27^. In particular, *B. subtilis* SUF includes two proteins that have been proposed as possible Fe-S scaffolds, SufB and SufU, whereas the *E. coli* SUF system includes only SufB. However, co-expressing *B. subtilis* SufU with *E. coli* SUF failed to activate additional orthologs (data not shown). Interestingly, all NadA orthologs that successfully complemented (with or without SUF expression) are phylogenetically grouped **(Figure 2A)**, implying that specific structural features might distinguish NadA orthologs that can be functionally expressed within *E. coli* from NadA orthologs inactive in *E. coli*. The NadA phylogeny is not congruent with known taxonomy (*i.e.* a mix of *Proteobacteria, Actinobacteria, Cyanobacteria* and *Archaea*, implying that the sources of NadA orthologs that are compatible with *E. coli* have a complex history of horizontal gene transfer.

We were surprised to find that electron carrier specificity may present a more stringent barrier to functional expression of IspG orthologs than the requirement for Fe-S cluster transfer. Although other redox-dependent enzyme classes (*e.g.* cytochrome P450 enzymes) are known to accept a limited range of electron carriers, many redox-dependent enzymes exhibit activity with multiple electron transfer proteins, and vice-versa. For instance, the *E. coli* flavodoxin FldA delivers electrons to BioB, IspG, and IspH^28^. Three *B. subtilis* electron transfer proteins (including YkuN and YkuP) are able to support activity of *B. subtilis* acyl lipid desaturase^13^ and five *Thermotoga maritima* ferredoxins are capable of supporting the rSAM enzyme MiaB^14^. Despite the fact that some redox enzymes might use electron transfer proteins interchangeably, prokaryotic genomes retain multiple electron transfer proteins that often show an extremely high level of sequence divergence, structural variation, and reduction potential^29^. The retention of multiple electron carriers indicates that each carrier is not compatible with all redox partners, despite some interchangability. We observed this selectivity as well: only one electron transfer protein from *R. sphaeroides* out of five tested activated *R. sphaeroides* IspG, and only two out of five *S. cattleya* electron transfer proteins activated *S. cattleya* IspG **(Table S6)**. Retaining an array of carriers enables species to support enzymes such as IspG or TsrM that specifically require low-potential electrons. In these cases, it is the need for low-potential reducing equivalents, rather than possession of an Fe-S cluster, that establishes a stringent barrier to functional gene transfer.

What do our results mean for using Fe-S enzymes in the context of engineered biosynthetic pathways? Overexpression of the host ISC and SUF pathways has been shown to improve the yield of properly-matured Fe-S enzymes^30^, likely because overexpression of Fe-S enzymes overtaxes the ability of the host Fe-S biogenesis systems to provide Fe-S clusters. However, we find that this approach may not reliably activate heterologous Fe-S enzymes. Instead, some may require co-expression with a heterologous Fe-S biogenesis pathway, as we observed for several NadA orthologs. If the Fe-S enzyme requires electron transfer, a specific electron transfer protein from the original host may be required. Given that prokaryotic genomes possess multiple electron carriers, identifying suitable electron transfer proteins may prove difficult. Suitable candidates may be identified based upon genome context of the Fe-S enzyme or by knowledge of its biochemical properties. In the case of TsrM, our awareness of the low reducing potential of cobalamin guided our selection to [4Fe-4S]-type ferredoxins. Some Fe-S enzymes require protein cofactors in addition to electron carriers. For instance, the rSAM enzyme lipoyl synthase (LipA) requires the *E. coli* Fe-S carrier protein NfuA to sustain catalytic turnovers^31^. In such cases, additional proteins from the original host may also be required to sustain heterologous Fe-S enzyme activity.

## Acknowledgements

We thank members of the Bokinsky Lab and the Barras Unit for discussions throughout the work, and Mohamed Atta and Julien Pérard for their assistance with biochemistry experiments. This project (IRONPLUGNPLAY) has received funding from the European Union’s Horizon 2020 research and innovation programme under grant 722361 and was supported by grants from Agence Nationale Recherche (ANR) ERA-Institut Pasteur and the ANR-10-LABX-62-IBEID. The work conducted by the U.S. Department of Energy Joint Genome Institute, a DOE Office of Science User Facility, is supported under Contract No. DE-AC02-05CH11231. SL was supported by the China Scholarship Council.

## Author Contributions

FD, EF, HS, and SL performed all cloning and complementation experiments. PG performed bioinformatics analysis. EF and RR engineered 2-methyltryptophan production. NvdB quantified 2-methyltryptophan by LCMS. F. Büke, CR, and M. Pabst measured expression of proteins by untargeted proteomics. M. Pelosse performed biochemical assays. ND performed electrochemistry experiments. BP, SG, SO-C, F. Barras, and GB supervised the project. All authors contributed to the final manuscript.

## Materials and Methods

### Selection of sequences and bioinformatics analysis

Orthologs of IspG, NadA, BioB, and ThiC selected in the preliminary screen were found in a variety of sequenced prokaryotic genomes using homology searches guided by *E. coli* sequences. To assemble the phylogenetically-representative distribution of NadA and IspG orthologs that was assembled following the preliminary screen, a systematic approach was used. First, a protein database was built from 248 representative prokaryotic proteomes **(Table S2)** (including 65 proteomes that were used for preliminary screen) gathered from NCBI FTP (ftp://ftp.ncbi.nlm.nih.gov/genomes/all), selecting around one genome *per* order (27 Archaea and 221 Bacteria) by using NCBI taxonomy and privileging complete proteomes. Homologues of NadA and IspG were first identified by BLASTp v2.8.1+^32^, using *E. coli* sequences as seeds (AAC73837.1 and AAC75568.1, respectively). Sequences associated to an *E*-value lower than 10^−4^ were aligned with MAFFT v7.419^33^ and used to build HMM profiles using hmmbuild from the HMMER v3.2.1 package^34^. These HMM profiles were then used to request the database using hmmsearch, and the sequences presenting an *E*-value lower than 10^−2^ were retrieved and aligned. Alignments were manually curated using Aliview v1.25^35^ and trimmed using BMGE v1.1^36^ with the substitution matrix BLOSUM30.

The best suited model for each alignment was selected using ModelFinder implemented in IQ-TREE v1.6.10^37^, according to the Bayesian Information Criteria (BIC) (LG+I+G4 for both NadA and IspG). Maximum Likelihood trees were inferred using PhyML v3.1^38^ and 100 bootstrap replicates performed to assess the robustness of the branches by using the transfer bootstrap value using BOOSTER v1.0^39^. For both NadA and IspG, the number of copies *per* genome was in the extreme majority equal to one. All homologues are thus considered as orthologues.

In order to select representative NadA and IspG sequences to be experimentally tested, the ML trees were split into clusters using TreeCluster v1.0^40^ with a threshold of 1.75, leading to 15 sequence clusters for NadA and 17 for IspG. Then, each cluster was further split into sub-clusters using hierarchical clustering implemented in scipy.cluster.hierarchy python library^41^. The number of sub-clusters for each cluster was set to 20% of the sequences within each cluster. One sequence per sub-cluster was finally selected, privileging reference strains and already tested sequences, leading to 47 sequences for NadA and 47 for IspG. The representativeness was checked by comparing the shape of distribution of patristic distances between the entire NadA and IspG trees and the selected sequences. The correlation between complementation data and phylogenetic distance was calculated by the correlation ratio *η*^2^. Data for phylogenetic analysis (containing protein database of the 248 prokaryotes, NadA/ IspG alignments and phylogenies and python script used to select sequences by clustering approach) is available at figshare.com/articles/dataset/Dangelo_et_al_Supplementary_data_zip/13664927.

### Bacterial strains, media and chemicals

The plasmids, oligonucleotides, and *E. coli* strains used in this study are listed in **Table S8**. *E. coli* Δ*ispG* was constructed by first integrating genes encoding the lower half of the mevalonate pathway to enable isoprenoid synthesis from exogenously-provided mevalonate. Genes encoding mevalonate kinase (MK), phosphomevalonate kinase (PMK) and mevalonate diphosphate decarboxylase (PMD) from *Saccharomyces cerevisiae* control by the P_trc_ promoter was amplified from pJBEI-2999^42^ and assembled by PCR to a FRT-flanked kanamycin resistance (KanR) cassette from pKD13 and integrated into *intA* using homologous recombination^43^. The KanR cassette was subsequently removed using the plasmid pCP20. *nadA, bioB,* and *thiC* were removed using homologous recombination in *E. coli* MG1655. Bacterial strains were routinely grown in aeration at 37°C in Luria-Bertani broth (LB) or in M9 medium (M9) supplemented with glucose (0.4%), CaCl_2_ (100 μM) and MgSO_4_ (1 mM). Solid media contained 1.5% agar. Anhydrotetracycline (aTc) 100 ng/mL, isopropyl β-D-1-thiogalactopyranoside (IPTG) 250 μM, mevalonate (MVA) 0.5 mM and nicotinic acid (NA) 10 μg/mL were added when indicated. When required, antibiotics were added at the concentrations of 50 μg/ml kanamycin (Kan) and 100 μg/mL ampicillin (Amp). Mevalolactone (MVL) and nicotinic acid (NA) were purchased from Sigma-Aldrich and resuspended in water at final concentrations of 1 M and 10 mg/mL, respectively. To prepare MVA, an equal volume of 1 M KOH was added to 1 M MVL and incubated at 37°C for 30 minutes.

### Plasmid construction for complementation assays

All the genes encoding heterologous orthologs and electron carrier proteins (with the exception of genes from *B. subtilis)* were codon-optimized for expression in *E. coli,* designed with strong ribosome binding sites, synthesized, and cloned into the expression vector pBbS2k^44^ by the Joint Genome Institute (JGI) directly, or were obtained as gene fragments by Twist Bioscience (TWB) **(Table S1 and S4)**, assembled, and cloned into pBbS2k. DH5α was used for all cloning steps. Genes encoding electron carrier proteins were codon-optimized for expression in *E. coli*, designed with strong ribosome binding sites, and assembled into the expression vector pBbA5a by JGI.

### Complementation assays

*E. coli* knockout strains were transformed with pBbS2k plasmid encoding corresponding orthologs and selected on permissive LB medium containing kanamycin. Plates supporting growth of *E. coli* Δ*ispG* strains additionally contained 0.5 mM MVA. For liquid culture-based complementation experiments, two individual colonies for each strain were inoculated into LB medium, grown for 7 hours at 37°C, and subsequently diluted 1:500 in M9 and added to M9 medium containing Kan and aTc. For reactivation experiments of inactive NadA orthologs, an *E. coli* Δ*nadA* strain bearing a plasmid encoding the *B. subtilis* SUF operon (pBbA5a-*sufCDSUB*) or the plasmid encoding the *E. coli* SUF operon (pBbA5a-*sufABCDSE*) was transformed with plasmids encoding orthologs and grown as described with the addition of 250 μM IPTG to induce expression of SUF (Table S6, S8). For reactivation experiments of inactive NadA orthologs, an *E. coli* Δ*ispG* strain bearing a plasmid encoding IspG orthologs was transformed with pBbA5a plasmids encoding electron transfer proteins. All growth curves have been performed in 96-wells microplates and cell density (OD_600_) was recorded by using an automated Spark 10Mluminometer-spectrophotometer (Tecan) for 18 hours (NadA experiments) or 12 hours (IspG experiments), every 30 minutes at 37°C in shaking condition. To test functionality of ThiC and BioB orthologs, *E. coli* Δ*bioB* and Δt*hiC* strains were transformed with pBbS2k plasmid encoding corresponding orthologs. Kanamycin resistant colonies were streaked on LB selective plates. The complementation test was performed by streaking on solid M9 glucose M9 medium supplemented with kanamycin and aTc. Plates were incubated at 37°C and growth was scored after 16 and 40 hours.

### SDS-PAGE test for protein expression

Δ*nadA* strains containing the pBbS2k plasmids and carrying NadA orthologs were grown in LB, Kan and aTc at 37°C to OD_600_ of 1.2 – 1.5. Δ*ispG* strains containing the pBbS2k plasmids carrying the IspG orthologs were grown in LB, Kan, MVA and aTc at 37°C to OD_600_ of 1.2 – 1.5. In both cases, the cells were centrifuged at 12,000 rpm for 2 min at room temperature and after removing the supernatant, the pellets were resuspended in PBS 1X buffer and then loaded on 12% SDS-PAGE gel after denaturating with Leammli loading dye at 100°C for 15 min. The gel was stained with Coomassie brilliant blue G-250 and decolorized using a solution of 60% water, 30% isopropanol, 10% acetic acid.

### Mass spectrometry detection of orthologs

#### Sample preparation

Δ*ispG* and Δ*nadA* strains bearing corresponding inactive orthologs were inoculated in 10 mL permissive media (Δ*ispG* in LB with 0.5 mM MVA, 100 μM IPTG, 25 ug/mL kanamycin; or Δ*nadA* in LB with 25 ug/mL kanamycin). When cultures reached early exponential phase (OD = 0.1), ortholog expression was maximally induced by 10 ng/mL doxycycline (added at early exponential phase before OD = 0.1). The cultures were subsequently grown to mid-logarithmic phase (OD = 0.4-0.6), at which point 2 mL solution of 10% (w/vol) trichloroacetic acid was added. Quenched samples were incubated on ice for at least 10 minutes before centrifugation (20,000 *g*, 4°C, 10 min), after which the supernatant was removed and pellet stored at −80°C before analysis.

#### Cell lysis and protein extraction

Briefly, biomass amount equivalent to 1 mL of 1 OD *E. coli* were collected in an Eppendorf tube and solubilised in a suspension solution consisting of 200μL B-PER reagent (Thermo Scientific) and 200μL TEAB buffer (50 mM TEAB, 1% (w/w) NaDOC, adjusted to pH 8.0) including 0.2μL protease inhibitor (P8215, Sigma Aldrich). Further, 0.1 g of glass beads (acid, washed, approx. 100 μm diameter) were added and cells were disrupted using 3 cycles of bead beating on a vortex for 30 seconds followed by cooling on ice for 30 seconds in-between cycles. In the following, a freeze/thaw step was performed by freezing the suspension at −80°C for 15 minutes and thawing under shaking at elevated temperature using an incubator. The cell debris was further pelleted by centrifugation using a bench top centrifuge at max speed, under cooling for 10 minutes. The supernatant was transferred to a new Eppendorf tube and kept at 4°C until further processed. Protein was precipitated by adding 1 volume of TCA (trichloroacetic acid, Sigma Aldrich) to 4 volumes of supernatant. The solution was incubated at 4°C for 10 minutes and further pelleted at 14,000 *g* for 10 minutes. The obtained protein precipitate was washed twice using 250μL ice cold acetone.

#### Proteolytic digestion

The protein pellet was dissolved to approx. 100 μg/100 μL of 200 mM ammonium bicarbonate containing 6 M Urea to a final concentration of approximately 100 μg/μL. To 100 μL protein solution, 30 μL of a 10 mM DTT solution were added and incubated at 37°C for 1 hour. In the following, 30 μL of a freshly prepared 20 mM IAA solution was added and incubated in the dark for 30 minutes. The solution was diluted to below 1 M urea using 200 mM bicarbonate buffer and an aliquot of approximately 25 μg protein were digested using sequencing grade Trypsin at 37°C overnight (Trypsin/Protein 1:50). Finally, protein digests were then further desalted using an Oasis HLB 96 well plate (Waters) according to the manufacturer protocols. The purified peptide eluate was dried using a speed vac concentrator.

#### One dimensional shot-gun proteomics approach

Briefly, the samples were analysed using a nano-liquid-chromatography system consisting of an ESAY nano LC 1200, equipped with an Acclaim PepMap RSLC RP C18 separation column (50 μm x 150 mm, 2 μm), and an QE plus Orbitrap mass spectrometer (Thermo). The flow rate was maintained at 350 nL/min over a linear gradient from 4% to 30% solvent B over 32.5 minutes, and finally to 70% B over 12.5 minutes. Solvent A was H_2_O containing 0.1% formic acid, and solvent B consisted of 80% acetonitrile in H_2_O and 0.1% formic acid. The Orbitrap was operated in data depended acquisition mode acquiring peptide signals form 385-1250 m/z at 70K resolution with a max IT of 100 ms and an AGC target of 3e6. The top 10 signals were isolated at a window of 2.0 m/z and fragmented using a NCE of 28. Fragments were acquired at 17K resolution with a max IT of 75 ms and an AGC target of 2e5. Singly charged, 6x and higher charged mass peaks were excluded from selection. Data were acquired from 0 to 60 min.

#### Database search

Data were analysed against the proteome database from *E. coli* K-12 (UniprotKB, TaxID 83333), including amino acid sequences for heterologous orthologs, using PEAKS Studio 10.0 (Bioinformatics Solutions Inc) allowing for 20 ppm parent ion and 0.02m/z fragment ion mass error, 2 missed cleavages, carbamidomethylation as fixed and methionine oxidation and N/Q deamidation as variable modifications. Peptide spectrum matches were filtered against 1% false discovery rate (FDR) and protein identifications with ≥ 2 unique peptides were accepted as significant.

### 2-methyl tryptophan biosynthesis

The mature protein coding sequence of the tryptophan 2-C-methyltransferase (TsrM) from *S. laurentii* (GenBank: FJ652572.1) was codon-optimized for *E. coli* expression and synthetized by Twist Bioscience. The optimized TsrM sequence was inserted into a pBbA5k BglBrick backbone between the BglII and BamHI sites to obtain the vector pTsrM. Codon-optimized genes encoding electron transfer proteins were cloned between the BamHI and XhoI sites of the pTsrM vector **(Table S8)**. The vector for cobalamin importer overexpression was constructed similarly to a previously-published report^23^. The genes for proteins BtuB (AYG21241.1, GenBank), BtuC (AYG19236.1, GenBank), BtuD (AYG19238.1, GenBank), BtuE (AYG19237.1, GenBank) and BtuF (AYG20683.1, GenBank) were amplified from *E. coli* MG1655 genome and cloned into the backbone pBbS2c BglBrick backbone to create the vector pBtu.

2-methyl tryptophan was produced using *E. coli* NCM3722 pBtu and with plasmids pTsrM, pTsrM_Scatt3Fd, pTsrM_Fdx1, pTsrM_Fdx2, or pTsrM_Fdx3. Individual colonies were picked and inoculated for pre-cultures grown overnight in 1 mL of SAM MOPS minimal medium (MOPS minimal medium^45^ supplemented with 1% w/vol glucose, 0.25% Casamino acids, and 7 μM hydroxocobalamin), at 37°C. The cultures (triplicates) were obtained by inoculating 50 μL of pre-cultures in 3 mL of SAM MOPS minimal medium. The cultures were grown at 37°C to OD_600_ of 0.05-0.1 for pBtu induction (83.3 ng/mL aTc) and continued to grow in the same conditions. At OD_600_ 0.5, TsrM and ferredoxin expression was induced (0.25 mM IPTG) and the medium supplemented with 0.15 mM cysteine and 32.5 μM FeCl_3_. Cultures were incubated at room temperature. Samples were taken after 24 hours (1 ml and the equivalent of 1 ml sample with 0.5 OD_600_ samples). Samples were centrifuged at 15,000 rpm for 2 min, and supernatants were transferred to new tubes. Pellets were quenched with NMM (methanol, acetonitrile and water, ratio of 5:3:1) + 0.1% formic acid and resuspended. Pellet samples were dried in speedvac at 40°C and resuspended with NMM for LC-MS measurement.

### 2-methyltryptophan quantification

2-methyl tryptophan was quantified by an LC-MS system (Agilent) using an XBridge BEH Amide 2.5 μm (Waters, Bridge Columns) with a precolumn and equipped with a standard ESI source mass spectrometer (sample injection volume of 5 μL). The mobile phase was compound by two solvents, A (20 mM ammonium formate in 10% acetonitrile) and B (20 mM ammonium formate in 80% acetonitrile). After 6 min at 100% solvent B, the metabolites were separated by a gradient from 100% to 70% of solvent B for 6 min (flow rate 0.4 mL/min), followed by a gradient from 70% to 100% for 50 sec (same flow rate) and held at 100% solvent B for 3 min 10 sec. The 2-methyl tryptophan precursor ion (positive polarity, 219.1) was fragmented into product ions (144.1 and 128) using an ESI ionization in MRM mode. 2-methyl tryptophan concentration was estimated using a calibration curve constructed with standard samples.

### Expression and purification of heterologous IspGs

Coding sequences of all tested IspG proteins were ordered codon optimized and cloned in a pET28a(+) plasmid by Genscript. A Tobacco etch virus (TEV) protease cleavage site was inserted between the N-Terminal His-tag and the downstream IspG sequences. Each IspG was expressed and purified under aerobic conditions. Expression was carried out in BL21DE3 cells for 3 hours under 1 mM IPTG induction at 37°C. Harvested cells were resuspended in lysis buffer (50 mM Tris pH 7.5, 250 mM NaCl, 10 mM imidazole), sonicated on ice (5 min, 10/30 sec ON/OFF, power 70%) and resulting lysate was clarified by ultracentrifugation (40,000 *g*, 20 min, 4°C). Supernatant was loaded onto a 5 mL Ni-NTA column. Column was washed using lysis buffer + 20 mM imidazole and proteins were eluted with lysis buffer + 300 mM imidazole. Fractions containing IspG were pooled together and incubated with TEV protease harbouring a His-tag (molar ratio: 1/100) o/n at 4°C for dialysis against storage buffer (50 mM Tris pH 7.5, 250 mM NaCl, 1 mM DTT). Dialysis bag content was reloaded onto 5 mL Ni-NTA column and flow-through (FT) containing IspG (without His-tag) were collected. Pure apo-IspG were concentrated and stored at −80°C. To counter instability of *E. coli* IspG, glycerol (50 % final) was added to concentrated purified protein which was subsequently stored at −20°C.

### Expression, purification and reconstitution of E. coli ErpA

*E. coli* ErpA was obtained as previously reported^46^.

### Fe-S cluster transfer between E. coli ErpA and heterologous IspGs

Fe-S cluster transfer experiment from *E. coli* ErpA to purified IspG orthologs was carried out in an anaerobic chamber (0_2_ < 2 ppm). For each transfer, one equivalent of apo-IspG (52 nmoles) from *E. coli*, *B. subtilis*, *S. cattleya, or Synechocystis sp. CACIAM05* was incubated in transfer buffer (50 mM Tris pH 7.5, 250 mM NaCl, 1 mM DTT) for 5 min before addition of 2.2 equivalents of [Fe-S]-ErpA from *E. coli* (115 nmoles). After 1 hour incubation, the mixture was loaded onto a 5 mL Ni-NTA column. Fractions containing [Fe-S]-IspG were collected in the FT whereas ErpA was eluted using elution buffer (50 mM Tris pH 7.5, 250 mM NaCl, 300 mM imidazole). Fractions were analysed by SDS-PAGE. UV-Vis spectra of ErpA and IspG were recorded before and after incubation and Fe-S transfer.

### TsrM overexpression and purification for in vitro studies

The coding sequence of TsrM from *S. laurentii* preceded by a N-terminal Tobacco etch virus (TEV) protease cleavable His-tag, was ordered codon optimized and sub-cloned into pET28a(+) vectors (Genscript). TsrM was co-expressed together with the Btu operon in BL21(DE3) cultured in LB medium supplemented with FeCl_3_ (50 μM), L-Cysteine (150 μM) and hydroxy-Cobalamin (2 μM). Btu operon expression was induced at OD_600_ = 0.3 using aTc (100 ng/mL) and TsrM expression was induced using IPTG (1 mM) at an OD_600_ = 0.7 for 18 hours at 18°C. Cells were harvested (6000 rpm, 20 min, 4°C), washed with NaCl 0.9 % and stored at −80°C. TsrM was purified purification under anaerobic conditions (0_2_ < 2 ppm) in a glove box (Jacomex). Cells expressing TsrM were resuspended in buffer containing 50 mM Tris pH 7.5, 250 mM NaCl, 10 mM imidazole, 10 % glycerol, 0.1 % Tween 20 and sonicated for 7 min (10/30 sec ON/OFF, power 50%). After ultracentrifugation (40,000 *g*, 20 min, 4°C), the soluble fraction was loaded onto a 5 mL Ni-NTA column equilibrated with buffer containing 50 mM Tris pH 7.5, 250 mM NaCl. After an extensive washing with buffer containing 50 mM Tris pH 7.5, 250 mM NaCl, 20 mM imidazole buffer, TsrM was eluted with buffer containing 50 mM Tris pH 7.5, 250 mM NaCl, 300 mM imidazole. Imidazole was removed using a HiPrep 26/10 desalting column equilibrated with buffer containing 50 mM Tris pH 7.5, 250 mM NaCl, 1 mM DTT. TsrM was then concentrated using an amicon cell until 37 mg/mL. As-isolated TsrM was then buffer exchanged into 25 mM Hepes pH 7.5, 300 mM KCl, 5% glycerol using micro Bio-spin 6 desalting column and used for electrochemistry (cyclic voltammetry) under anaerobic conditions or aliquoted for storage in liquid N_2_.

### Expression and purification of *S. cattleya* ferredoxins

Coding sequences of the three ferredoxins from *S. cattleya* (AEW96347.1 (Fdx1), AEW92689.1 (Fdx2), AEW97532.1 (Fdx3)), all preceded by a N-terminal Tobacco etch virus (TEV) protease cleavable His-tag, were ordered codon optimized for expression in *E. coli* and sub-cloned into pET28a(+) vectors (Genscript). Each ferredoxin was expressed in BL21(DE3) cells cultured in LB medium. Protein expression was induced using IPTG (1 mM) for 3 hours at 37°C. Cells were harvested (6000 rpm, 20 min, 4°C) and washed with NaCl 0.9 % before storage at −80°C. All ferredoxins were purified under anaerobic conditions (0_2_ < 2 ppm) in a glove box. Cells expressing ferredoxins were resuspended in buffer containing 50 mM Tris pH 7.5, 250 mM NaCl, 10 mM imidazole, 10 % glycerol, 0.1 % Tween 20 and sonicated for 7 min (10/30 sec ON/OFF, power 50%). After ultracentrifugation (40,000 *g*, 20 min, 4°C), the soluble fractions were loaded onto 3 separate 5 mL Ni-NTA columns equilibrated with 50 mM Tris pH 7.5, 250 mM NaCl. Columns were washed using buffer containing 50 mM Tris pH 7.5, 250 mM NaCl, 20 mM imidazole and proteins were eluted with buffer containing 50 mM Tris pH 7.5, 250 mM NaCl, 300 mM imidazole. Imidazole was removed using a HiPrep 26/10 desalting column, equilibrated with a buffer containing 50 mM Tris pH 7.5, 250 mM NaCl, 1 mM DTT. Ferredoxins cleared of imidazole were incubated overnight in presence of TEV (molar ratio: 1/100). The mixtures were reloaded onto 5 different 5 mL Ni-NTA columns and flow-through (FT), containing ferredoxins, were collected.

### Fe-S reconstitution of *S. cattleya* Fdx1

Fdx2 and Fdx3 from *S. cattleya* were purified anaerobically with their Fe-S clusters in contrast to Fdx1 which was obtained as an apo-form. Fdx1 was therefore reconstituted within an anaerobic chamber. Fdx1 was incubated in buffer containing 50 mM Tris pH 7.5, 250 mM NaCl, 1 mM DTT with 10-fold equivalents of Fe^2+^ (Mohr salt) and 10-fold equivalents of S^2−^(Na_2_S). The reaction occurred for 3 hours at RT and after centrifugation (15 min at 16,000 *g*) the mixture was loaded onto a Superdex-75 10/300 column. Fractions containing reconstituted Fdx1 were pooled and concentrated using concentrators for microfuge. Subsequent analysis of Fe content and concentration determination were performed as described below.

### Biochemical analyses of TsrM

Concentration of proteins (TsrM and ferredoxins) was determined using Rose Bengal with BSA as standard^47^ and the Fe content was determined using the Fish method^48^. The cobalamin content of as-isolated TsrM was determined by UV-Vis absorption spectroscopy through its conversion to dicyanocobalamin (ɛ_367_ = 30,800 M^−1^cm^−1^) using a treatment with 0.1 M potassium cyanide following a procedure previously reported^21^.

### Protein-Film Electrochemistry

Protein-film electrochemistry experiments were performed anaerobically inside an anaerobic chamber (O_2_ < 2 ppm) using freshly purified TsrM, [Fe-S]-containing ferredoxins and a potentiostat (Biologic). A three-electrode configuration was used in a small volume analytical cell (Biologic) with a platinum wire and an Ag/AgCl electrode as counter and reference electrode respectively. When analysing TsrM, a pyrolytic graphite edge (PGE) electrode was used to collect electrochemical measurements. The electrode was polished with sand paper 1200 followed by 1 μm alumina, and then baseline measurements were collected by placing the PGE electrode into the buffer cell solution (10 mM MES, 10 mM CHES, 10 mM TAPS, 10 mM HEPES pH 8, 200 mM NaCl). Then, 3 μL of 350 μM TsrM were applied to the polished PGE electrode for 5 min before being rinsed with 200 μL of the buffer cell solution. Next, the electrode was immediately placed back in the buffer cell solution for measurements. For ferredoxins, a glassy carbon (GC) electrode polished using 1 μm alumina was used within a setup combining a small volume analytical cell and the sample holder of the SVC-2 kit (Biologic). Ferredoxin solutions were at 50 μM (in 10 mM MES, 10 mM CHES, 10 mM TAPS, 10 mM HEPES pH 8, 200 mM NaCl buffer). Cyclic voltammograms were collected at room temperature with a scan rate of 100 mV/sec. Redox potentials of TsrM and ferredoxins were determined through square wave voltammetry (SWV) under the same conditions as previously described for cyclic voltammetry. The SWV input signal consisted of a staircase ramp from −1 to −0.3 V vs. Ag/AgCl, with 2 mV increments, 50 mV stair amplitude and 5 Hz frequency. Electrochemical signals were analysed by correction of the non-Faradaic component from the raw data using the QSoas package^49^.

### TsrM-ferredoxin affinity measurements

Affinity interactions between TsrM and *S. cattleya* ferredoxins were measured by BioLayer Interferometry inside an anaerobic chamber (O_2_ < 2 ppm) equipped with a BLItz system (FortéBio). All proteins were buffer exchanged into 25 mM Hepes pH 7.5, 300 mM KCl, 5% glycerol using micro Bio-spin6 desalting columns. Ferredoxins were biotinylated through incubation with 1 equivalent of NHS-PEG_4_-Biotin. After 30 min, excess NHS-PEG_4_-Biotin was removed using a micro Bio-spin 6 desalting column. Biotinylated ferredoxins (10 μg/mL) were bound for 60 sec to streptavidin biosensors equilibrated in BLI buffer (25mM Hepes pH 7.5, 300 mM KCl, 5% glycerol) containing Tween 20 and BSA to inhibit non-specific binding, following manufacturer recommendations. After having been plunged for 20 sec for equilibration in BLI buffer, the loaded-biosensor was plunged into TsrM solution for 80 sec, followed by 60 sec into BLI buffer for association constant and dissociation constant measurements, respectively. This procedure was repeated for various concentrations of TsrM. Affinity constants were extracted from fitted raw data using the BLItz software (PALL/FortéBio).

## Notes

### Competing Interest Statement

The authors have declared no competing interest.

