## Supplemental Figures & Text for "Resolving phylogenetic and biochemical barriers to functional expression of heterologous iron-sulphur cluster enzymes"

**Affiliations:** [a] Department of Bionanoscience, Kavli Institute of Nanoscience, Delft University of Technology, Delft, The Netherlands; [b] Department of Microbiology, UMR Institut Pasteur-CNRS 2001, Unit Evolutionary Biology of the Microbial Cell, Paris, France; [c] Department of Microbiology, UMR Institut Pasteur-CNRS 2001, Unit Stress Adaptation and Metabolism of Enterobacteria, Paris, France; [d] Univ. Grenoble Alpes, CNRS, CEA, IRIG, Laboratoire de Chimie et Biologie des Métaux, Grenoble, France; [e] Aix-Marseille Université-CNRS, Laboratoire de Chimie Bactérienne, Institut de Microbiologie de la Méditerranée, Institut Microbiologie Bioenergies Biotechnologie, Marseille, France; [f] Department of Biotechnology, Delft University of Technology, Delft, The Netherlands

**Current addresses:** [1] Institute for Translational Vaccinology (Intravacc), Process Development Bacterial Vaccines, Bilthoven, The Netherlands; [2] EMBL Grenoble, Grenoble, France; [3] Instituto de Tecnologia Química e Biológica António Xavier, Universidade Nova de Lisboa, Oeiras, Portugal; [4] Vaccine Process & Analytical Development, Janssen Vaccines & Prevention B.V., Leiden, The Netherlands

### Supplemental Figures

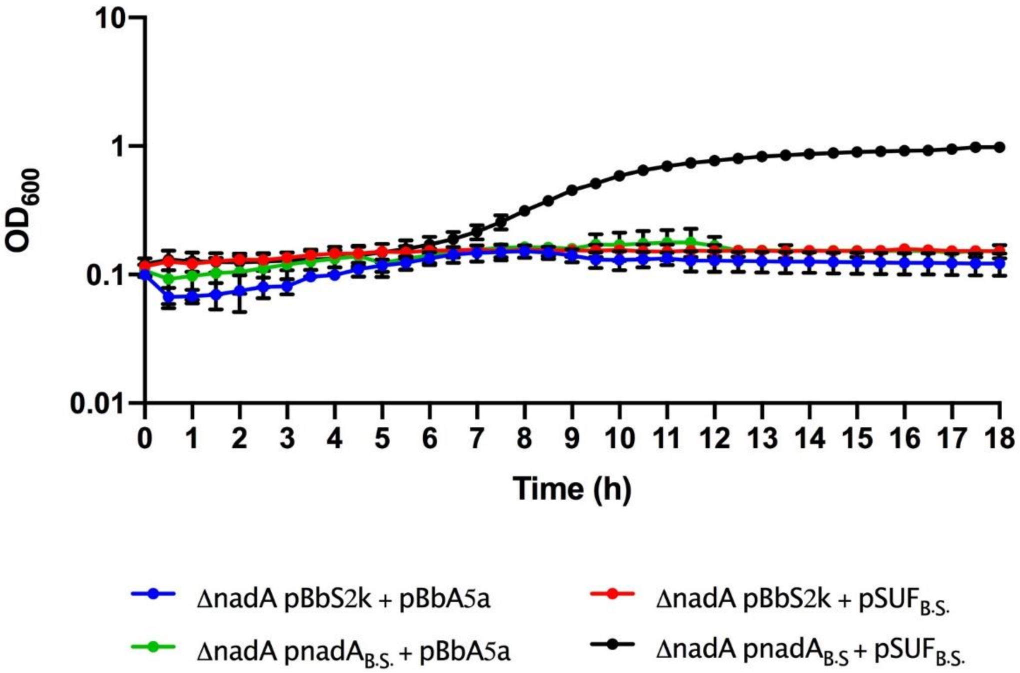


**Figure S1.** Growth curves of *E. coli* Δ*nadA* grown in M9, Kan, Amp, aTc and IPTG complemented with pBbS2k and pBbA5a plasmids carrying different genes: empty vectors (blue line), pBbS2k and *B. subtilis* SUF (red line), *B. subtilis nadA* and pBbA5a (green line) and *B. subtilis nadA* and *B. subtilis* SUF (black line). *B.S.*: *B. subtilis.* The average of two biological replicates is reported with SD.

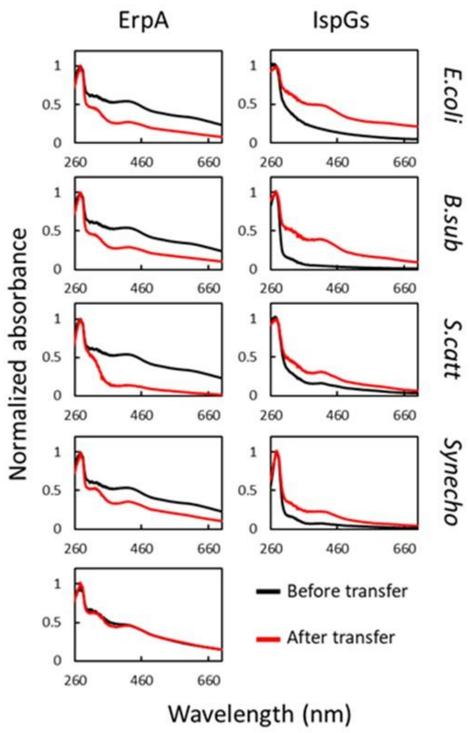


**Figure S2.** *E. coli* ErpA mediates Fe-S cluster transfer into heterologous IspGs. UV-visible spectra of [Fe-S]-ErpA and apo-IspGs were recorded before ErpA/IspG co-incubation (black traces). One equivalent of apo-IspG was incubated with 2.2 equivalents of [Fe-S]-ErpA. After 1 h incubation, proteins were separated on an affinity column (IspGs in the flow-through (FT), ErpA in the eluate). Spectra were recorded for both (red traces). The bottom left panel corresponds to spectra of *E. coli* ErpA incubated without IspG. IspG orthologs used: *E. coli*: *E. coli*; *B. sub*: *B. subtilis*; *S. catt: S. cattleya; Synecho: Synechocystis sp. CACIAM05.*

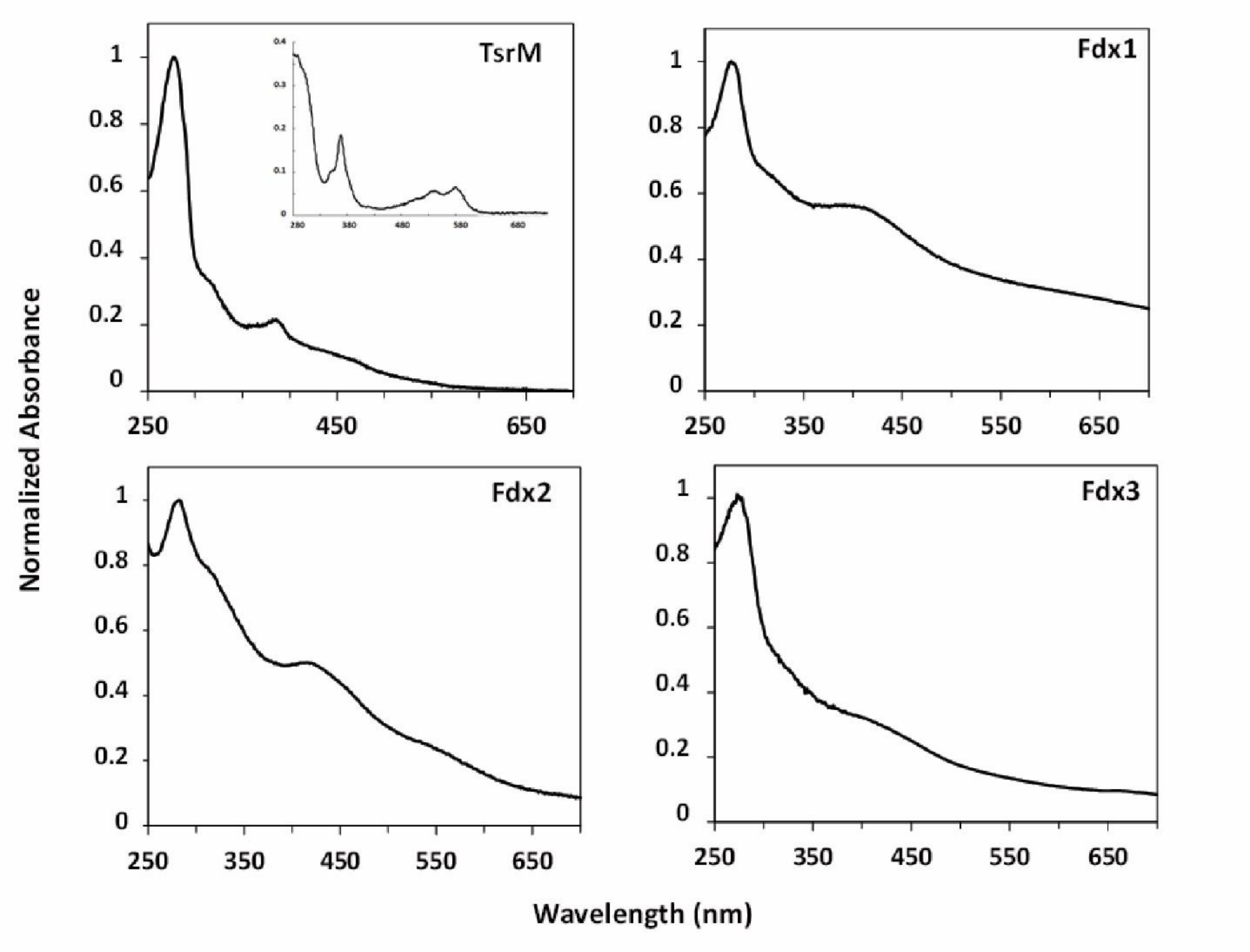


**Figure S3.** TsrM, Fdx1, Fdx2 and Fdx3 with their metallic cofactors. Protein concentration were normalized with $Abs_{280} = 1$. Consistent with previous observations, as-purified TsrM (0.9 Cbl and 3.5 Fe ± 0.3 per monomer) shows features consistent with the presence both of Fe-S cluster and of cobalamin, including the broad absorption between 350 and 600 nm, indicative of a Fe-S cluster, and the distinct feature at 390 nm, suggestive of the presence of cob(I)alamin. Inset: TsrM treated with potassium cyanide yields the dicyanocobalamin adduct of cobalamin, which has a strong feature at 367 nm ($\varepsilon_{367}$ = 30,800 $M^{-1}$ $cm^{-1}$). Concerning *S. cattleya* ferredoxins (Fdx1, Fdx2, Fdx3), their iron content was consistent with their annotations as di-cluster and single-cluster ferredoxins (7.4 Fe ± 0.5/monomer, 2.9 Fe± 0.1 /monomer, and 2.8 Fe± 0.1 /monomer for Fdx1, Fdx2, and Fdx3, respectively).

### Supplementary Tables

Due to their size, the Supplementary Tables are provided as tabular data files.

<Table S1.csv>

**Table S1**. Accession numbers, nucleotide sequences of orthologs of NadA, IspG, BioB, ThiC, and IspD used in the preliminary screen. Complementation results incidated with minus (-): negative complementation; plus (+): positive complementation; (+/-): growth detected but substantially slower than with *E. coli* ortholog; N.D.: not determined.

<Table S2.csv>

**Table S2.** List of genomes used for the phylogenetic analysis.

<Table S3.csv>

**Table S3**. List of the NadA and IspG orthologs retrieved from the database depicted in Table S2. Sequences that have been selected for the complementation test are indicated by a (+) in the adjacent column.

<Table S4.csv>

**Table S4.** Accession numbers and nucleotide sequences used to create a phylogenetically-representative selection of NadA and IspG orthologs.

<Table S5.csv>

**Table S5**. Results of SDS-PAGE and shotgun proteomics of selected orthologs. Orthologs detected by visible bands on the gel are indicated by a (+).

<Table S6.csv>

**Table S6**. List of the Fe-S biogenesis proteins and electron carriers used to test re-activation of IspG orthologs using complementation. Re-activation of IspG orthologs was tested first by co-expressing all genes as a multi-cistronic operon. For cases in which individual electron carriers and Fe-S biogenesis proteins were sub-cloned for complementation testing, proteins identified to activate heterologous IspG orthologs (positive complementation) are indicated by (+), while proteins that did not activate IspG (no complementation) are indicated by (-).

<Table S7.csv>

**Table S7**. Results of the IspG complementation and reactivation experiments using electron carrier proteins listed in Table S6. Minus (-): negative complementation; plus (+): positive complementation; (+/-): growth detected but substantially slower than with *E. coli* ortholog; N.D.: not determined. The rightmost column, provided to summarize reactivation experiments, indicates IspG orthologs that were successfully activated by at least one electron carrier attempted.

<Table S8.csv>

**Table S8**. List of the bacterial strains, plasmids, and oligonucleotides used in this study.
